# Detection of Enterovirus D68 in wastewater samples from the United Kingdom during outbreaks reported globally between 2015 and 2018

**DOI:** 10.1101/738948

**Authors:** Manasi Majumdar, Thomas Wilton, Yara Hajarha, Dimitra Klapsa, Javier Martin

**Affiliations:** Division of Virology, National Institute for Biological Standards and Control (NIBSC), South Mimms, Potters Bar, Herts EN6 3QG, UK

**Keywords:** wastewater, human enterovirus, acute flaccid myelitis, enterovirus d68, sewage, environmental surveillance

## Abstract

Detection of enterovirus D68 (EV-D68) in wastewater samples from the UK between December 2014 and December 2018 showed a marked seasonal distribution with a high proportion of samples containing EV-D68 during periods when identification of this virus in clinical samples was most common. This includes a recent upsurge of EV-D68 detection in respiratory samples from the United Kingdom between August and December 2018 associated with cases of acute flaccid myelitis, following similar reports in the USA. Phylogenetic analysis of EV-D68 sewage strains demonstrated that strains belonging to distinct genetic clades followed the same temporal distribution as that observed for EV-D68 clinical strains in the UK and that they showed very close genetic relationship with EV-D68 strains circulating elsewhere in the world during the same periods. The results demonstrated a clear association between detecting EV-D68 in wastewater and finding it in clinical samples which was somehow unexpected given that EV-D68 is rarely detected in stool samples. We conclude that the use of environmental surveillance is a valuable tool to detect and monitor outbreaks due to EV-68 infection.

## INTRODUCTION

Infection with enterovirus D68 (EV-D68) has been associated with severe respiratory disease in humans with increasing evidence of its link to neurological complications causing acute flaccid myelitis (AFM), a polio-like syndrome resulting in long-term or permanent disability (Cassidy et al. 2018). Although EV-D68 was first identified in 1962, it was rarely reported until 2008 when small outbreaks started to emerge in what appears to be mostly a biannual distribution (Kramer et al. 2018). The first known large outbreak of severe respiratory illness associated with EV-D68 infection occurred in the USA between August and December 2014 (Holm-Hansen et al. 2016). Genetically related EV-D68 strains were also found during the same period in Canada, Europe and Asia with more than 2,000 cases reported in 20 countries (Kramer et al. 2018). These outbreaks were temporally and geographically associated with an increase in AFM cases, particularly in the USA, and a similar association was observed in Europe, Argentina and the USA in 2016 (Cassidy et al. 2018). An increase in AFM cases was again reported in the USA in 2018 with 230 confirmed cases mostly between August and November (Centers for Disease Control and Prevention (CDC) 2019). Concurrently, a sharp increase of EV-D68 detections in clinical samples from respiratory cases was detected which also peaked in September 2018 (Kujawski et al. 2019). The Centres for Disease Control in the USA concluded that the clinical, laboratory, and epidemiologic evidence suggested a clear viral association with AFM cases although not necessarily with EV-D68 alone (McKay et al. 2018). Similarly, a sharp rise in polio-like AFM cases has also been seen in the UK with 40 AFM cases reported in England in 2018 (The United Kingdom Acute Flaccid Paralysis Afp Task 2019), the majority of which occurring since September 2018. Fifteen cases were associated with enterovirus; EV-D68 was detected in 9, enterovirus C104 in 1, coxsackievirus B1 in 1, rhinovirus in 1 and, in 3 cases, the enterovirus was not typeable (The United Kingdom Acute Flaccid Paralysis Afp Task 2019).

In order to investigate the circulation patterns of EV-D68 in the United Kingdom, we analysed a total of 118 wastewater samples taken between December 2014 and December 2018 for the presence of EV-D68. The results were compared to those reported from clinical samples both in terms of date of detection and genetic signature.

## METHODS

Sewage specimens have been collected monthly in Glasgow and London since December 2014 and June 2016, respectively. Samples up to December 2018 were processed using WHO standard protocols (Majumdar et al. 2017) and analysed for the presence of EV-D68 using recently established methods (Majumdar et al. 2018). Briefly, RT-PCR products containing coding sequences for EV-D68 capsid VP1 protein were generated using a nested PCR protocol. Whole-capsid Pan-Enterovirus RT-PCR products were first synthesized directly from RNA extracted from sewage concentrates and then used as templates for a second PCR reaction targeting the genomic region coding for EV-D68 capsid protein VP1 as described before (Majumdar et al. 2018). Previously described VP1 EV-D68 universal primers VP1-F (5’-ACCATTTACATGCRGCAGAGG-3’) and 485 (5’-ACATCTGAYTGCCARTCYAC-3’) were used (Imamura et al. 2013, Oberste et al. 2004). Additional in-house primers specific for EV-D68 genetic clades (sequences available on request) were utilized and hence sequences from different genogroups were obtained from the same sample in some cases. Purified VP1 PCR products were sequenced by the Sanger method with an ABI Prism 3130 genetic analyser. EV-D68 VP1 sequences obtained in this study were compared to EV-D68 sequences available in the GenBank database to establish geographical and temporal phylogenetic relationships between them. The Molecular Evolutionary Genetics Analysis (MEGA) software package version 7.0 (Kumar et al., 2016) was used for these analyses. The phylogenetic tree was drawn using FigTree version 1.4.2 program (http://tree.bio.ed.ac.uk/software/figtree/). Relevant VP1 nucleotide sequences from this study have been deposited in GenBank (NCBI Accession Numbers MK377389-377406). Statistical analyses to test the association between detection of EV-D68 in wastewater and clinical samples were performed using Fisher’s exact test.

## RESULTS AND DISCUSSION

As shown in Fig. 1, 21 out of the 118 wastewater samples analysed were positive for EV-D68 showing a marked seasonal distribution. EV-D68 was identified in wastewater samples during four separate periods: Dec-2014 to Jan-2015, Sep-2015 to Jan-2016, Jun-2016 to Sep 2016 and Aug-2018 to Dec-2018 with an isolated sample from Glasgow being positive in September 2017 (Fig. 1). We used weekly EV-D68 clinical detection data available from Wales (Cottrell et al. 2018) to compare them with environmental surveillance (ES) results in London and Glasgow during the same calendar week and found a statistically significant association between the probability of finding EV-D68 in sewage and detecting it in clinical samples (Fig. 1). A high specificity and positive/negative predictive values for detecting EV-D68 in wastewater samples as compared to finding it in clinical cases in Wales were observed but sensitivity for EV-D68 sewage detection was relatively low (62.5 and 40.0% for London and Glasgow, respectively). This is not entirely unexpected as wastewater samples were only collected on a single day monthly and the volume of sewage tested by PCR was low (equivalent to 5ml of raw sewage). Nevertheless, EV-D68 was frequently detected in wastewater samples during reported outbreaks despite the fact that the virus is rarely found in stools from clinical cases (Barnadas et al. 2017, Van Leer-Buter et al. 2016). Although the presence of EV-D68 in sewage has been reported before (Benschop et al. 2017, Brinkman et al. 2017, Weil et al. 2017), our finding was unexpected and suggests that EV-D68 might replicate in the gut more commonly than it is thought, perhaps during the initial stages of infection before respiratory symptoms develop and samples are collected for analysis. Although unlikely, the possibility that EV-D68 present in human respiratory fluids could reach the sewage system through washing, showering, etc. cannot be ruled out.

**Figure 1.**
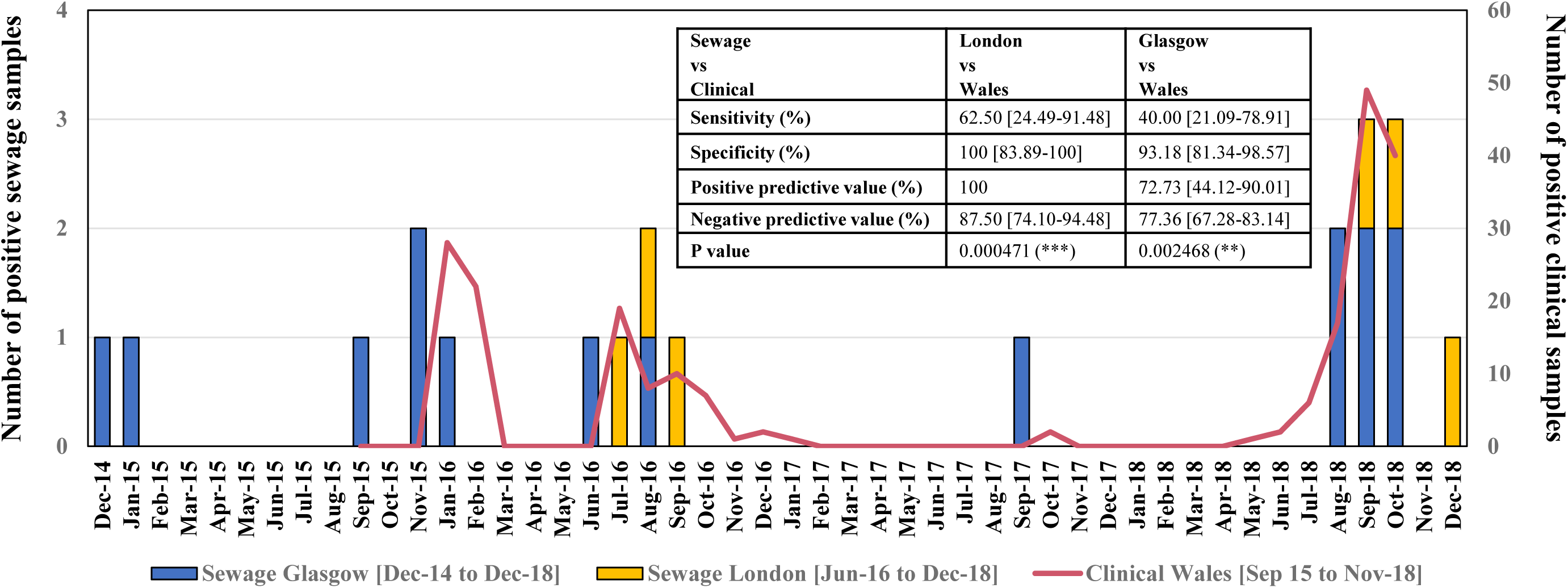
Identification of EV-D68 in wastewater samples from the UK. The presence of EV-D68 in sewage was analyzed using a PanEV+VP1 nested PCR assay followed by Sanger sequencing (Majumdar et al. 2018).The number of monthly wastewater samples positive for EV-D68 are shown in blue (Glasgow) or orange (London) columns. EV-D68 clinical data from Wales, covering the period between 1 September 2015 to 5 November 2018, are shown as a red line (results according to Cottrell et al, 2018 (Cottrell et al. 2018)). Monthly samples from Dalmarnock and Shieldhall (Glasgow) were taken from 17 December 2014 to 27 December 2018 with the exception of December 2015 in Dalmarnock and December 2015, April-June 2016, September-November 2016 and July 2017 in Shieldhall when samples were not taken for technical reasons. Monthly samples from Beckton (London) were taken from 14 June 2016 to 13 December 2018. The inlet Table shows results of statistical analysis to test the association between detection of EV-D68 in wastewater and clinical samples performed using Fisher’s exact test and comparing weekly clinical data from Wales with sewage data from London and Glasgow using actual collection dates to assign the corresponding week.

Statistical association between wastewater and clinical EV-D68 detection data was better when using London (located closer to Wales) sewage data than data from Glasgow which suggests minor regional variations in virus transmission. As shown in Fig 1, detection of EV-D68 in the Glasgow sewage appeared to have preceded that in London sewage and clinical samples from Wales during the 2015-2016 outbreaks. Wastewater samples from Glasgow collected on 25 September and 25 November 2015 were positive for EV-D68, seven and two weeks before the first cases were reported in Wales, respectively (Fig. 1). Similarly, EV-D68 was detected in a wastewater sample from 29 June 2016 in Glasgow only few days before EV-D68 clinical cases were reported (Cottrell et al. 2018). In addition, a single wastewater sample from 25 Sep 2017 was positive for EV-D68, four weeks before the only two clinical cases found between the 2016 and 2018 outbreaks were reported in Wales, suggesting that ES can also detect low levels of virus transmission. During the 2018 outbreak, few clinical cases positive for EV-D68 were reported in Wales before the virus was first found in sewage (Cottrell et al. 2018). Nevertheless, detection of EV-D68 in wastewater and clinical samples largely overlapped. Clinical detection of EV-D68 in London during the same periods followed very similar temporal distribution (Public Health England 2019) although data were not available for detailed statistical comparisons.

The first EV-D68 season detected in sewage corresponded to the last stages of a period during which large numbers of EV-D68 strains associated with respiratory disease were reported worldwide in what was the first known large EV-D68 outbreak (Holm-Hansen et al. 2016). Following this event, identification of EV-D68 in wastewater samples showed very close resemblance to clinical data reported in the UK, where peaks of EV-D68 detection in human respiratory samples were observed in January 2016, July 2016 and September 2018, and very few cases were reported between November 2016 and July 2018, as shown in the two separate studies from Wales and England discussed above (Cottrell et al. 2018, Public Health England 2019). Reports of EV-D68 detection were also abundant in other European countries during the same periods in 2016 and 2018 (Bal et al. 2019, Cassidy et al. 2018). An upsurge of EV-D68 detection in wastewater samples from both locations was observed between August and December 2018 in agreement with recent reports of increased EV-D68 detection likely associated with AFM cases in the UK, following similar reports in the USA (Centers for Disease Control and Prevention (CDC) 2019, The United Kingdom Acute Flaccid Paralysis Afp Task 2019).

Phylogenetic analysis of the UK EV-D68 sewage strains (Fig. 2) confirmed that wastewater samples contained an accurate representation of EV-D68 genetic clades identified in clinical samples in the UK. Although very few actual nucleotide sequences from EV-D68 clinical strains from the UK are publicly available, information on the relative abundance of EV-D68 strains from different genetic clades present in clinical samples from England between 2014-2018 has been reported (Public Health England 2019). The temporal distribution of EV-D68 genetic clades among EV-D68 strains identified in clinical samples closely resembled that of the EV-D68 strains found in wastewater samples as determined from the genetic analysis described here. Very similar genetic clade distribution of EV-D68 clinical strains was observed in Europe and North America although EV-D68 clinical and sewage detections in the UK did not appear to follow a clear-cut biannual distribution as reported elsewhere. Nevertheless, most UK EV-D68 sewage strains showed very close genetic relationship (>99% VP1 sequence similarity) with clinical strains from Europe and USA found during the same periods, in many cases showing identical or nearly identical VP1 sequences to sequences from contemporary viruses available from GenBank. EV-D68 strains from genetic clade D and sub-clade B2, found both in clinical and wastewater samples in the UK at the end of 2014 and beginning of 2015 showed very close genetic relationship to viruses from France, Germany and Italy (Fig. 2). In particular, the two sewage strains from December 2014 and January 15 were genetically very close to clinical strains KP657742 (NCBI ID) and KP745743 (99.77% and 99.77% VP1 sequence identity, respectively), both found in Germany in October 2014. Sub-clade B3 viruses were detected in clinical and wastewater samples from the UK during the 2015 and 2016 outbreaks. All EV-D68 sewage strains during these two outbreak periods belonged to genetic subclade B3 and showed very close genetic relationship between them and with contemporary clinical strains found globally showing very high VP1 sequence similarity with them (Fig. 2); e.g. with strain LC107892 from Japan in August 2015 (99.77%) and many clinical strains identified across Europe (Denmark, France, Germany, Italy, Sweden and The Netherlands) and the USA between April and September 2016 with VP1 sequence similarity as high as 99.76-100%. The single strain found in the Glasgow sewage in 2017 belonged to genetic clade D as did the very few clinical strains identified during that period in England (Public Health England 2019). This virus was genetically related to a clinical strain found in the USA also in September 2017 (MG757146; 99.03% VP1 sequence identity). Active co-circulation of EV-D68 strains from both genetic clades D and sub-clade B3 was observed between August and December 2018 in the UK as strains from these two genotypes were found in wastewater and clinical samples. EV-D68 strains from both genetic clades D and sub-clade B3 were genetically very close to 2018 clinical strains found in Sweden, France and Italy (Fig. 2). For example, a sub-clade B3 virus identified in the Glasgow sewage on 29 June 2016 (MN018239) contained a highly similar VP1 sequence to viruses found in clinical samples from New York (KY385889; 99.77%, June 2016), Sweden (MH674122; 99.77%, August 2016) and Germany (KX830909, 99.66% July 2016). Similarly, a clade D strain found in the London sewage on 11 September 2018 (MN018255) contained identical VP1 sequence to a virus strain identified in a clinical sample taken on 24 September 2018 in Italy (MK301345). Only one B3 strain (MN018253), found in a wastewater sample collected in October 2018 in London, was somehow different to all other B3 sewage strains showing a more distant genetic link to clinical strains from India; e.g. strains MH733832 from July 2017 and MH330334 from September 2017, with 98.55% and 97.82% VP1 sequence identity, respectively. This indicates the possible presence of a minor variant in the population.

**Figure 2.**
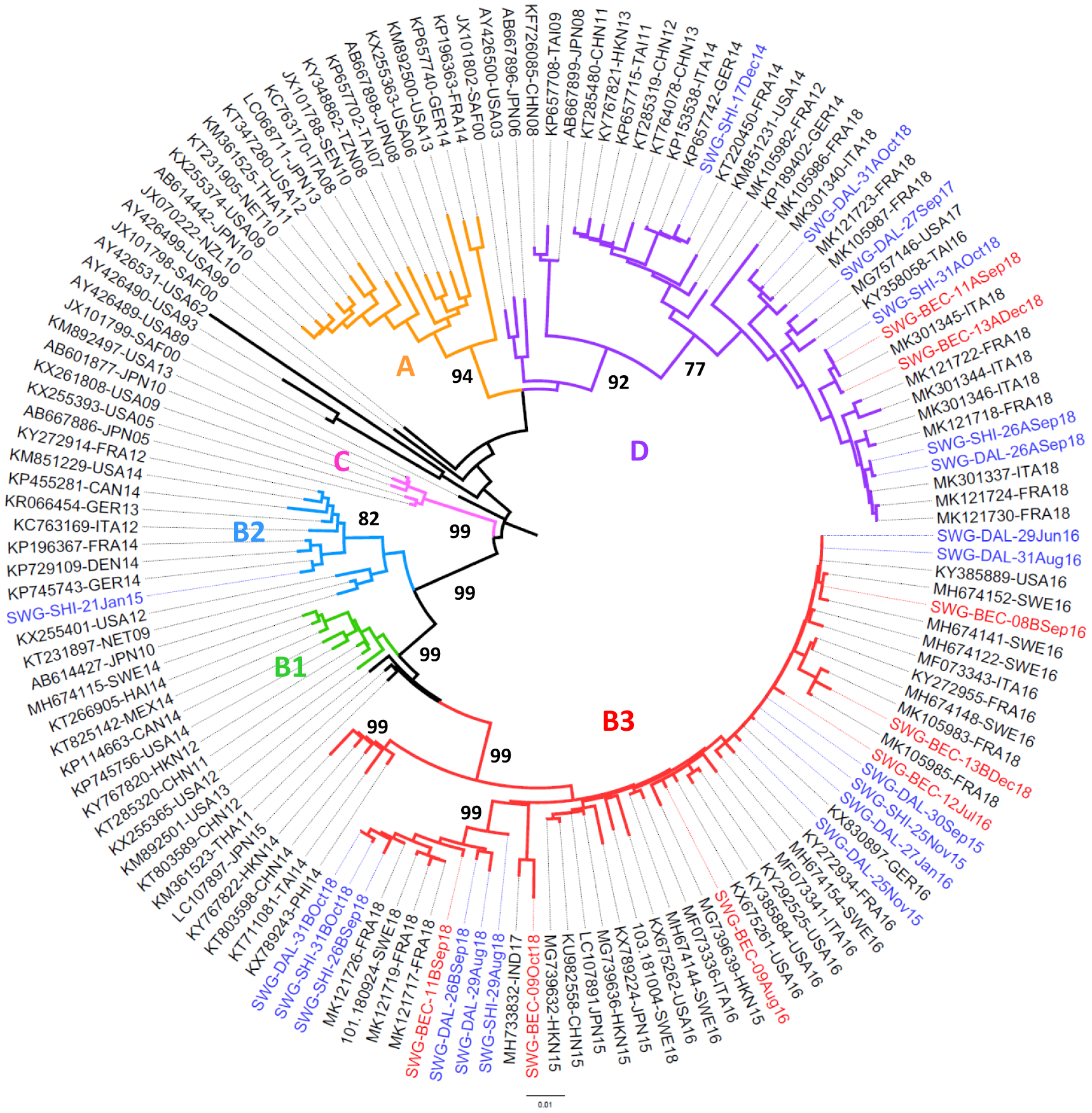
Phylogenetic analysis of EV-D68 strains found in UK wastewater samples. The evolutionary histories of EV-D68 strains were inferred using the Neighbor-Joining method with VP1 sequences of the sewage strains and representatives of all EV-D68 genetic clades from GenBank. Sequences for 2018 strains from Sweden were downloaded from http://virological.org/t/enterovirus-d68-3-sequences-from-sweden-2018/258. The evolutionary distances were computed using the Maximum Composite Likelihood method expressed in base substitutions per site. The optimal tree is shown with Bootstrap values (1000 replicates) indicated next to relevant branches. Names for phylogenetic groups (clades and sub-clades) are indicated (A, B1, B2, B3 and D). EV-D68 sewage strains from the UK are shown in the format SWG-XXX-YyyZZ in which XXX represents location (DAL=Dalmarnock, SHI=Shieldhall and BEC=Beckton), Yyy represents month and ZZ represents year with viruses shown in red (London) or blue (Glasgow). Viruses indicated with an A or a B mean indicate two different variants identified in that particular sample. Accession number, location and year of isolation for GenBank sequences are indicated in the sequence names. Abbreviations for country names are ITA: Italy; USA: United States of America; MEX: Mexico; JPN: Japan; CHN: China; THA: Thailand; GER: Germany; SAF: South Africa; NZL: New Zealand; TAI: Taiwan; NET: Netherlands; SEN: Senegal; TZN: Tanzania; FRA: France; DEN: Denmark; CAN: Canada; PHI: Philippines; HKN: Hong Kong; SWE: Sweden; IND: India; HAI: Haiti.

The high sequence similarity observed between strains found in London and Glasgow wastewater samples and between these strains and clinical strains found globally suggest a very rapid and widespread transmission of EV-D68 during outbreaks, most likely due to its respiratory transmission pathway. Thus, importation events might contribute to the initiation of outbreaks. It is likely that EV-D68 transmission mainly occurs during rapidly transmitting short-term outbreaks and that increased association of EV-D68 infection with severe disease might be a consequence of increased virus transmission and not necessarily due to virus mutations conferring increased virulence. Nevertheless, thorough genotypic and phenotypic analysis of EV-D68 strains found between 1962 and now would be required to further assess if significant changes in EV-D68 genotype/phenotype are responsible for the observed increase in transmission and disease severity associated with EV-D68 infection.

Our results indicate that ES did not appear to detect silent transmission of EV-D68 as no positive ES samples were identified in periods when there were no clinical cases reported. Hence, it is not clear how much EV-D68 silent transmission occurs between outbreak periods, although the absence of EV-D68 detection in clinical and wastewater samples during these long periods suggests little transmission. Apart from few exceptions of EV-D68 detections in the Glasgow sewage, detection of EV-D68 in wastewater samples did not precede finding it in clinical samples, which means our ES system, as currently set up, will not be an adequate alert system for EV-D68 outbreak detection. This is likely due to low sensitivity of EV-D68 detection in sewage as discussed above, given the limited sampling conducted. However, sensitivity for EV-D68 detection could potentially be improved by increasing the frequency of sampling and the volume of sewage analysed as it has been shown for poliovirus. ES has proven to be a very sensitive system to detect poliovirus circulation even in the absence of paralytic disease(Asghar et al. 2014). The current recommendation is to collect samples at least twice monthly and to test a minimum of concentrated sewage volume equivalent to 150 ml of raw sewage. ES for EV-D68 could also be very useful as a supplementary surveillance system to clinical surveillance to identify gaps and to certify interruption of circulation during outbreaks. This will be particularly useful in areas were surveillance for EV-D68 is poor or non-existent. ES samples can be further analysed for the presence of other enterovirus serotypes with possible implication in human disease (Majumdar et al. 2018).

## Conclusions

- Direct detection of EV-D68 in sewage concentrates using our nested PCR system is a highly sensitive method for the detection of circulating EV-D68 causing outbreaks despite the virus being rarely found in stool samples from clinical cases.
- The temporal distribution of EV-D68 genetic clades found in wastewater samples closely resembled that seen in clinical samples with EV-D68 sewage strains showing very close genetic similarly to EV-D68 clinical strains found globally.
- Environmental surveillance for EV-D68 can be used as a supplementary surveillance system to monitor outbreaks and to evaluate the quality of clinical surveillance.

## ACKNOWLEDGEMENTS

We would like to thank Dr Catherine Moore and Dr Simon Cottrell from Public Health Wales for providing published clinical data from Wales. This report is independent research that, in part, was funded by the Department of Health Policy Research Programme (NIBSC Regulatory Science Research Unit, 044/0069) and the Bill and Melinda Gates Foundation (Funder ID: 100000865) (Opportunity/Contract ID: OPP1171890). All authors declare no conflict of interests.

